# Urine proteomic profiling at admission reveals complement biomarkers linked to alcohol-associated liver disease

**DOI:** 10.64898/2026.04.04.716339

**Authors:** Luan Prado, Ryan Musich, Moyinoluwa Taiwo, Vai Pathak, Daniel M. Rotroff, Annette Bellar, Nicole Welch, Jaividhya Dasarathy, David Streem, the AlcHepNet, Srinivasan Dasarathy, Laura E. Nagy

**Author notes:** Address correspondence to: Laura E. Nagy, Cleveland Clinic, Lerner Research Institute/NE40 9500 Euclid Ave, Cleveland OH 44195. DMR reports research funding from Bayer Pharmaceuticals and Novo Nordisk; a license agreement with OpenDNA for diabetes stratification; consulting fees and equity in Genovation Health LLC and ClarifiedPrecision Medicine; and is an inventor on patents related to liver cancer biomarkers, precision oncology decision support, and GLP-1RA responsiveness (International Patent Application No. PCT/US2022/28611 and US Provisional Patent Application No. 63/737,152). The other authors have declared that no conflict of interest exists. This work was supported in part by NIH grants: P50AA024333 (to LEN, SD and DMR), R01AA030699 (to LEN), U01AA026398 (to LEN), K08AA028794 (to NW), R01GM119174 (to SD), R01DK113196 (to SD), R01AA021890 (to SD), U01AA026976 (to SD), R56HL141744 (to SD), U01DK061732 (to SD), U01DK062470 (to SD), See Supplemental Acknowledgments for details on the AlcHepNet Consortium. The timsTof Pro2 instrument was purchased via an NIH shared instrument grant, S10 OD030398.

## Abstract

**Background and aims:** Circulating complement is associated with occurrence of alcohol-associated hepatitis (AH) and is a potential biomarker to distinguish AH from alcohol cirrhosis (AC). Complement contributes to kidney injury, a condition often occurring in patients with alcohol-associated liver disease (ALD). However, little is known regarding complement in cross talk between liver and kidney in ALD. Here we tested the hypothesis that urinary complement would provide potential biomarkers for ALD and insights into mechanisms of liver-kidney crosstalk in the pathogenesis of ALD.

**Methods:** Plasma and urine were collected at admission from patients with sAH, healthy controls (HC), and heavy drinkers without liver disease (HD) (from the multicenter Alcohol Hepatitis Network) and with AC (from the Northern Ohio Alcohol Center). Urine was subjected to unbiased proteomics analysis and plasma complement assessed by multiplex/ELISA assays. 30- and 90-day mortality was tracked in patients with sAH.

**Results:** All three complement activation pathways were perturbed in plasma and urine of patients with sAH and AC compared to HC and HD. Components of the lectin and classical pathways in urine were associated with 30- and 90-day mortality in patients with sAH. When 4 complement proteins were combined, they distinguished sAH from AC (AUC 0.78), equivalent to that of MELD (AUC 0.65). There was no correlation between complement in plasma and urine, suggesting an independent impact of sAH on complement in kidney and liver.

**Conclusion:** The urinary proteome revealed complement protein signatures associated with sAH and AC, providing valuable insights into the potential for complement biomarkers and the mechanisms of liver-kidney crosstalk in ALD.

## Introduction

Alcohol-associated liver disease (ALD) encompasses a range of conditions resulting from chronic, heavy alcohol use. Alcohol-associated hepatitis (AH) is an acute condition marked by a rapid development of severe inflammation in the liver. While AH can present in individuals with heavy daily alcohol intake and no underlying ALD, individuals with underlying fibrosis or alcohol cirrhosis (AC), a chronic liver disease characterized by ongoing inflammation and fibrosis ^1,2^, are at higher risk of developing AH^3,4^. Given that AC be an underlying ALD in patients with sAH, there is a need to identify new biomarkers for sAH or AC that can reliably differentiate these conditions and better predict patients with AC likely to develop sAH.

In patients with AH or AC, there is strong activation of innate immune responses. It is well established that cellular components of the innate immune system, including tissue-resident macrophages and peripheral monocytes and neutrophils recruited to the liver, produce multiple cytokines and chemokines that drive the persistent inflammation seen in AH and contribute to inflammation and fibrosis in AC ^3,5–8^. However, much less is known about the role of complement in the progression of AH and AC ^7,9^. Complement, part of the innate immune system, is responsible for ongoing immune surveillance. Complement is activated through three pathways: classical, alternative, and lectin. Although each pathway is initiated via different signals, all three converge to form the C3- and C5-convertases, which generate the anaphylatoxins, C3a and C5a, and lead to the formation of the membrane attack complex (MAC) ^10–14^. Specific complement components can either enhance or negatively regulate complement activation and function. Enhancers, such as Factor D (CFD), are particularly important for the alternative pathway ^14–17^. Negative regulators prevent excessive activation of complement and protect host cells from injury and death. These include membrane-bound proteins such as CD55, CD59, and CD46 ^18–20^, as well as soluble regulators such as factor H (CFH), related proteins (CFHR1–5) ^21,22^, and factor I (CFI) ^23^. Additionally, proteins such as uromodulin (Umod, also known as Tamm-Horsfall protein), though not exclusive to the complement system, play a key role in regulating and fine-tuning complement activation ^24,25^.

Recent studies have revealed extensive perturbations in the expression of both complement activators and negative regulators in the circulation and liver of patients with AH ^9,26^. Significantly, the concentration of the anaphylatoxin C5a is increased in the plasma of patients with severe AH (sAH), suggesting that the net impact of complement dysregulation results in complement activation ^26^. Interestingly, multiple complement components in the circulation, including collectin 11 (CL11), C1q binding protein (C1qBP), CD55, and CD59, can distinguish patients with sAH from healthy controls and individuals with alcohol use disorder (AUD). More clinically relevant, circulating concentrations of C1qBP, complement component 8 (C8), CD55, and CD59 can accurately distinguish patients with sAH from those with AC ^9^, suggesting that complement could contribute to the non-resolving inflammation characteristic of sAH, and could be developed as an informative diagnostic biomarker.

Complement is also known to contribute to kidney injury due to a variety of etiologies. However, while kidney injury is a condition often occurring in patients with ALD, very little is known regarding the contribution of complement in crosstalk between the liver and kidney in sAH or AC. Here we utilized an unbiased proteomics approach to discover that urinary complement signatures can differentiate patients with sAH from heavy drinkers without liver disease and from patients with AC. Independent perturbations in plasma and urinary complement also indicate important liver-kidney cross talk related to complement in patients with sAH.

## Methods

### Study populations

Samples from three different clinical studies were used in this study: 1) the multicenter Alcohol-associated Hepatitis Network (AlcHepNet) Observational study (NCT 3850899) ^27,28^; 2) the prednisolone control arm of the AlcHepNet Randomized Clinical Trial (RCT) (NCT04072822), a multicenter prospective clinical investigation ^29^; and 3) the Northern Ohio Alcohol Center (NTC03224949). This study was approved by the Institutional Review Boards of all participants’ institutions, and all participants signed written consent before data collection and urine and blood sample collection. Both male and female patients were included in all cohorts studied here.

#### Cohort 1 – plasma complement cohort

Demographic and clinical data of the subjects used in the plasma complement cohort are summarized in Table 1. All samples were collected during the first visit after enrollment (i.e., baseline sampling). Complement components were quantified in plasma from 156 subjects enrolled in the AlcHepNet Observational study, including 106 patients with severe AH (sAH), 25 heavy drinkers without liver disease (HD), and 25 healthy controls (HC). In the sAH group, 25 of these samples overlapped with samples in the urinary proteomics cohort. The samples included in this analysis were based on the availability at the time of the assay.

**Table 1.** Cohort 1 – Plasma complement cohort: clinical and demographic data of the patients.

|  | HC (N=25) | HD (N=25) | Severe AH (N=106) | p value |
| --- | --- | --- | --- | --- |
| **Sex** |  |  |  | ns |
| **Female** | 13 (52.0%) | 10 (40.0%) | 33 (31.1%) |  |
| **Male** | 12 (48.0%) | 15 (60.0%) | 73 (68.9%) |  |
| **Race** |  |  |  | <0.0001 |
| **American Indian/Alaska Native** | 0 (0.0%) | 1 (4.0%) | 0 (0.0%) |  |
| **Asian** | 5 (20.0%) | 0 (0.0%) | 0 (0.0%) |  |
| **Black/African American** | 3 (12.0%) | 3 (12.0%) | 8 (7.5%) |  |
| **Unknown** | 0 (0.0%) | 0 (0.0%) | 2 (1.9%) |  |
| **White** | 17 (68.0%) | 21 (84.0%) | 96 (90.6%) |  |
| **Age (years) (Mean± SD)** | 45.24±10.87 | 43.84±10.79 | 45.04±9.98 | ns |
| **MELD (Mean± SD)** | 6.68±0.99<sup>a</sup> | 6.48±1.39<sup>a</sup> | 29.97±7.53<sup>b</sup> | <0.0001 |
| **CPS (Mean± SD)** | 5.00±0.00<sup>a</sup> | 5.16±0.47<sup>a</sup> | 9.85±1.97<sup>b</sup> |  |
| **Total Bilirubin (mg/dL) (Mean± SD)** | 0.53±0.25<sup>a</sup> | 0.46±0.25<sup>a</sup> | 21.89±11.86<sup>b</sup> | <0.0001 |
| **Serum creatinine (mg/dL) (Mean± SD)** | 0.86±0.14<sup>a</sup> | 0.83±0.14<sup>a</sup> | 2.00±1.91<sup>b</sup> | 0.0031 |
| **eGFR (mL/min/1.73m<sup>2</sup>) (Mean± SD)** | 87.20±16.27<sup>ab</sup> | 95.76±24.96<sup>b</sup> | 74.65±59.97<sup>a</sup> | 0.0056 |
| **BMI (Mean± SD)** | 26.88±4.82<sup>a</sup> | 28.12±6.70<sup>ab</sup> | 30.44±6.40<sup>b</sup> | 0.0082 |
| **INR (Mean± SD)** | 1.02±0.11<sup>a</sup> | 1.02±0.14<sup>a</sup> | 1.98±0.55<sup>b</sup> | <0.0001 |
| **Albumin (g/L) (Mean± SD)** | 4.36±0.20<sup>a</sup> | 4.26±0.46<sup>a</sup> | 2.97±0.63<sup>b</sup> | <0.0001 |
| **ALT (U/L) (Mean± SD)** | 20.84±11.92<sup>a</sup> | 29.28±16.87<sup>a</sup> | 57.95±42.29<sup>b</sup> | <0.0001 |
| **AST (U/L) (Mean± SD)** | 22.28±10.18<sup>a</sup> | 29.44±15.33<sup>a</sup> | 136.26±89.35<sup>b</sup> | <0.0001 |

3 One-way ANOVA followed by least square means for normal distributed variables and Kruskal-Wallis test followed by Dwass, Steel, 4 Critchlow-Fligner test non-normal distributed variables. Fisher's exact test for the categorical variables. Means followed by different 5 letters differ between themselves. MELD: model of end-stage liver disease; CPS: Child-Pugh Score; INR: International normalized 6 ratio; ALT: alanine aminotransferase; AST aspartate aminotransferase; eGFR: estimated glomerular filtration rate; ns: non- 7 significant.

#### Cohort 2 – urinary proteomics

A total of 134 subjects were included (Table 2). Patients with sAH were followed for 30- and 90-day mortality. De-identified urine samples, along with clinical and demographic data, were obtained from 27 HC, 25 HD, and 27 patients with sAH from the AlcHepNet Observational study, and 34 patients with sAH from the AlcHepNet RCT prednisolone arm. For the AC group, 21 patients diagnosed with AC from the Northern Ohio Alcohol Center were included. All samples were collected during the first visit after enrollment (i.e., baseline urine sampling) for patients enrolled in the AlcHepNet studies. For patients with AC, samples were collected during routine check-ups as part of disease follow-up.

**Table 2.** Cohort 2 – Urine proteomics cohort: Clinical and demographic data of the patients.

|  | HC (N=27) | HD (N=25) | AC (N=21) | sAH (N=61) | p value |
| --- | --- | --- | --- | --- | --- |
| **Sex** |  |  |  |  | ns |
| **Female** | 13 (48.1%) | 10 (40.0%) | 7 (33.3%) | 19 (31.1%) |  |
| **Male** | 14 (51.9%) | 15 (60.0%) | 14 (66.7%) | 42 (68.9%) |  |
| **Age (years) (Mean±SD)** | 46.07±10.99<sup>a</sup> | 47.92±13.63<sup>a</sup> | 51.19±10.80<sup>a</sup> | 43.43±9.18<sup>b</sup> | <0.0001 |
| **Race** |  |  |  |  | 0.0021 |
| **Asian** | 5 (18.5%) | 0 (0.0%) | 0 (0.0%) | 1 (1.6%) |  |
| **Black/African American** | 3 (11.1%) | 6 (24.0%) | 4 (19.0%) | 2 (3.3%) |  |
| **Unknown** | 0 (0.0%) | 0 (0.0%) | 0 (0.0%) | 1 (1.6%) |  |
| **White** | 19 (70.4%) | 19 (76.0%) | 17 (81.0%) | 57 (93.4%) |  |
| **MELD (Mean±SD)** | 6.78±1.01<sup>a</sup> | 6.68±1.55<sup>a</sup> | 16.78±8.67<sup>a</sup> | 29.21±8.20<sup>b</sup> | <0.0001 |
| **CPS (Mean±SD)** | 5.00±0.00<sup>a</sup> | 5.12±0.44<sup>a</sup> | - | 7.71±2.94<sup>b</sup> | 0.0035 |
| **Total Bilirubin (mg/dL) (Mean±SD)** | 0.56±0.25<sup>a</sup> | 0.45±0.17<sup>a</sup> | 4.17±5.3<sup>b</sup> | 24.21±11.7<sup>c</sup> | <0.0001 |
| **BMI (Mean±SD)** | 27.13±4.73<sup>a</sup> | 28.10±6.99<sup>a</sup> | 25.99±4.61<sup>a</sup> | 30.95±6.96<sup>b</sup> | 0.0145 |
| **WBC (x10<sup>3</sup>) (Mean±SD)** | 5.91±1.33<sup>a</sup> | 6.47±2.13<sup>a</sup> | - | 13.04±7.95<sup>b</sup> | <0.0001 |
| **INR (Mean±SD)** | 1.03±0.11<sup>a</sup> | 0.99±0.15<sup>a</sup> | 1.44±0.41<sup>b</sup> | 1.97±0.52<sup>c</sup> | <0.0001 |
| **Albumin (g/L) (Mean±SD)** | 4.37±0.20<sup>a</sup> | 4.27±0.39<sup>a</sup> | 3.11±1.02<sup>b</sup> | 2.95±0.65<sup>b</sup> | <0.0001 |
| **Serum Creatinine (mg/dL) (Mean±SD)** | 0.87±0.14 | 0.85±0.21 | 1.43±1.67 | 1.83±1.74 | ns |
| **eGFR (mL/min/1.73m<sup>2</sup>) (Mean±SD)** | 79.59±13.45<sup>a</sup> | 76.44±19.81<sup>a</sup> | - | 60.69±36.49<sup>b</sup> | 0.0015 |
| **ALT (U/L) (Mean±SD)** | 21.19±11.88<sup>a</sup> | 26.12±17.21<sup>a</sup> | 25.29±12.47<sup>a</sup> | 52.18±36.87<sup>b</sup> | <0.0001 |
| **AST (U/L) (Mean±SD)** | 22.22±9.90<sup>a</sup> | 23.92±7.76<sup>a</sup> | 55.14±37.71<sup>b</sup> | 138.58±81.75<sup>c</sup> | <0.0001 |
| **Total Protein (g/L) (Mean±SD)** | 7.18±0.31<sup>a</sup> | 7.03±0.63<sup>a</sup> | - | 5.74±0.85<sup>b</sup> | <0.0001 |

2 One-way ANOVA followed by least square means for normal distributed variables and Kruskal-Wallis test followed by Dwass, Steel, 3 Critchlow-Fligner test non-normal distributed variables. Fisher's exact test for the categorical variables. Means followed by different 4 letters differ between themselves. MELD: model of end-stage liver disease; CPS: Child-Pugh Score; INR: International normalized 5 ratio; ALT: alanine aminotransferase; AST aspartate aminotransferase; eGFR: estimated glomerular filtration rate; ns: non- 6 significant.

### Plasma Complement

Complement components C1q, C3, C4, C4b, C5, CFB, CFD, CFI, CFH, and iC3b were quantified using multiplex assays by Exsera BioLabs in Aurora, Colorado, USA (https://www.exserabiolabs.org). The concentrations of complement soluble C5b9 (MicroVue^TM^ SC5b-9 Plus EIA, Quidel, A020) and CL11 (Human Collectin-11, MyBioSource, MBS2883570) were measured by ELISA.

### Proteomics

The method was developed in-house by the Cleveland Clinic Proteomics and Metabolomics Core based on published methods ^30–33^. Briefly, urine samples were centrifuged at 15000 x g for 20 minutes at room temperature (RT), and the supernatant was filtered through a 0.5 mL Amicon 3 kD MWCO filter. Then, the samples were centrifuged at 14000 ×g for 15 minutes. Samples were incubated in acetone at −20 °C overnight, then pelleted and air-dried. Samples were reconstituted in 50 µL of urea lysis buffer (100 mM Tris-HCl buffer, 8 M urea, 1X protease inhibitor cocktail, pH 8.0).

For protein quantification, 2 µL from each sample was taken for the BCA assay, and 20µg of protein from each sample was used for in-solution digestion. Samples were reduced by dithiothreitol, alkylated by iodoacetamide, and precipitated using −20°C acetone overnight. Samples were then pelleted and air-dried. The dried samples were re-suspended in 40 µL 50 mM triethyl ammonium bicarbonate (TEAB) and digested using 0.5 µg trypsin per sample. After overnight incubation, the digested samples were centrifuged at 21,000 x g for 15 minutes. Four micrograms of the digest from each sample were then transferred to a new tube, spun using a SpeedVac, and reconstituted in 16 µL of 0.1% formic acid. Finally, a 10 µL sample was mixed with 10 µL of 0.5x iRT standards and analyzed by LC-MS using data-independent acquisition (DIA). Four µL of the sample was injected, and the peptides eluted from the column (BurkerFifteen 15 cm x 75 µm) by an acetonitrile/0.1% formic acid gradient at a flow rate of 0.3 µL/min. Then the eluate was injected into the LC-MS system (Bruker timsTOF pro2 mass spectrometry system interfaced with a Bruker NanoElute HPLC system).

### Data processing and statistical analyses

For categorical variables, Fisher’s exact test was performed. For the multiplex and ELISA analysis, data were log-transformed and compared using ANOVA, followed by a Bonferroni post-hoc analysis. The ANOVA p-values were corrected using False Discovery Rate (FDR), and FDR adjusted p-value <0.1 were considered significant.

In the standard differential expression analysis of the urinary proteome, all samples were evaluated by PCA to investigate potential batch effects. There was no clear separation between diagnoses or between batches (Supplemental Figure 1A and B). Relative abundance values from LCMS were used. Missing values were replaced with 0s, and 0.001 was added to all counts to avoid “divide by 0” errors for fold change calculations. Volcano plots and PCA plots were created in R (v4.5.0) using *ggplot2* (v3.5.2).

For individual comparisons for urinary complement, data were log-transformed. Comparison among the four groups was performed using ANOVA, followed by Bonferroni post-hoc analysis. The p-values were FDR-adjusted, and a q < 0.1 was considered significant. Pairwise comparison of the urine complement of patients who died and those who did not was performed using Student’s t-test. Continuous variables are presented as Mean±SD.

For the ROC curves^34^ (Figure 4A, C, D) protein abundance was normalized using a log2 transformation. Individual protein discriminatory performance was assessed by ROC analysis, with 95% confidence intervals calculated by the DeLong method ^35^. Multi-protein panel performance (Figure 4 B) was assessed using logistic regression with leave-one-out cross-validation (LOOCV) in SAS (SAS Institute, version 8.2.5 1277), with center-and-scale preprocessing applied within each fold to prevent data leakage. AUC and 95% confidence intervals are reported for each pairwise comparison.

All statistical analyses were done using SAS or R version 4.5.0 ^36^. Graphs were made using GraphPad Prism 10 (version 10.0.2).

## Results

### Plasma complement is dysregulated in patients with sAH but not heavy drinkers

Expression of complement proteins is dysregulated in the plasma of patients with sAH and AC ^9,26^. However, it is not fully understood if these changes are a consequence of heavy alcohol consumption, liver disease, or a combination of both ^9^. Therefore, using multiplex and ELISA assays, we compared plasma complement concentrations in healthy controls, heavy drinkers, and patients with sAH (Figure 1). FDR-adjusted p-values for each protein are summarized in the supplemental data (Supplemental Table 1). Of all the analytes measured, there were no differences in the concentrations between HC and HD (Figure 1). However, the concentrations of multiple complement components differed in plasma from patients with sAH compared with both HC and HD. In the alternative pathway, CFB and CFD, involved in activation of the alternative pathway, were upregulated while CFI and CFH, negative regulators of the pathway, were downregulated in plasma of patients with sAH compared to HC and HD (Figure 1A). In the classical pathway, no difference was observed in C1q, while C4 and its activation fragment C4b were down regulated in plasma of patients with sAH compared to HC and HD (Figure 1B). The concentration of CL11 was increased in plasma of patients with sAH compared to HC and HD (Figure 1C). The two proteins common to all three activation pathways, C3 and C5, did not differ between groups (Figure 1D). Notably, the complement activation fragment, iC3b, was increased in sAH compared to both HC and HD, indicating that the net impact of changes in complement in sAH tends to promote complement activation. Interestingly, sC5b9, the soluble form of the MAC, was not different in the plasma of patients with sAH compared with HC and HD (Figure 1E). Overall, these findings suggest that dysregulation of the complement is linked to sAH rather than to heavy drinking.

**Figure 1.**
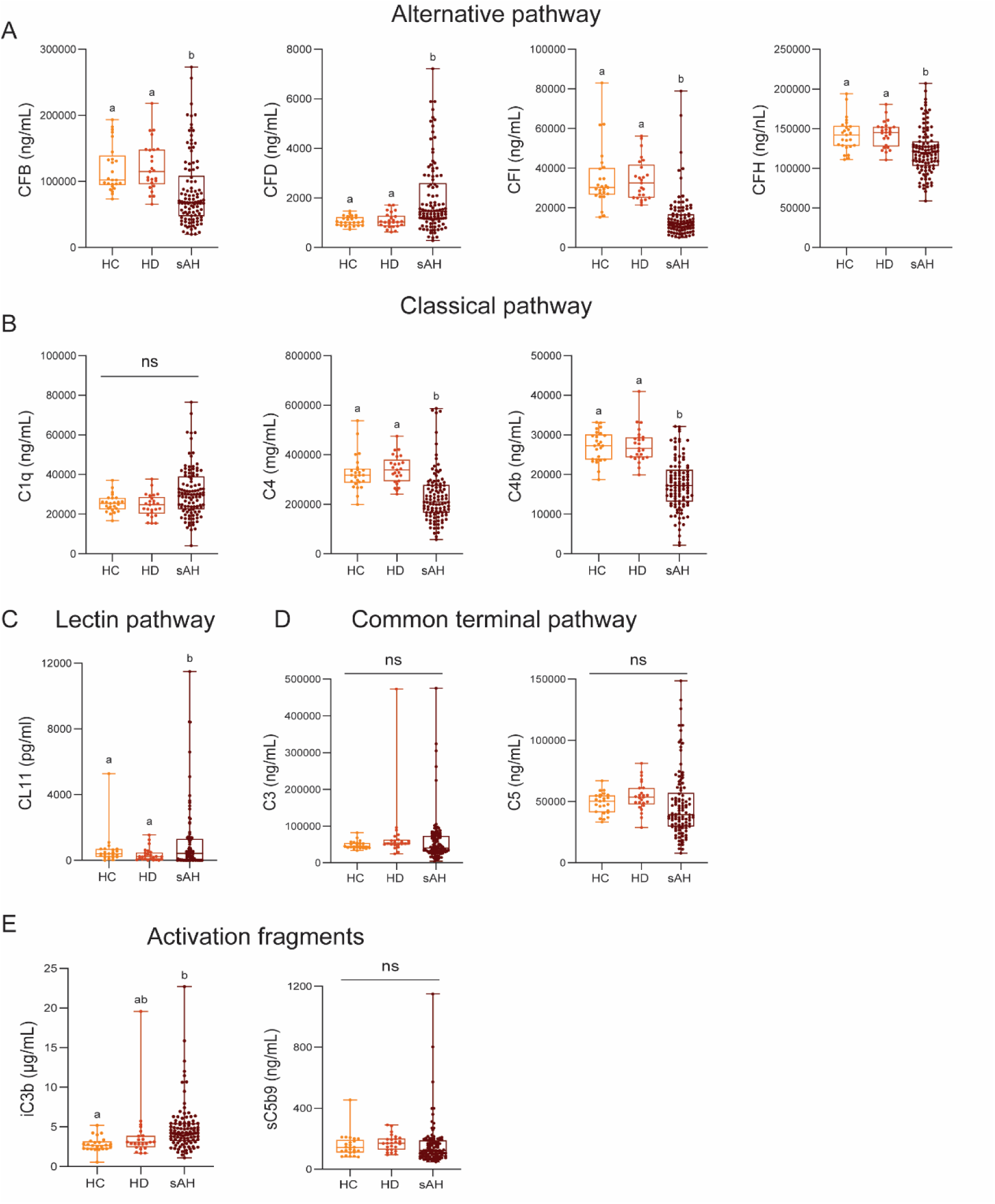
Plasma complement is perturbed and activated in patients with sAH from cohort 1. A) Alternative pathway B) Classical pathway C) CL11 of the lectin pathway. D) Complement C3 and C5, from the common terminal pathway. E) Activation fragments iC3b and sC5b9. Box-and-whisker plots display minimum and maximum values, lower and upper quartiles, and the median, with each individual value represented as a point superimposed on the graph. ANOVA with FDR-adjusted p-value (q<0.1), followed by Bonferroni post hoc test for individual group comparisons. Different letters mean significant differences between groups’ mean values.

### Characterization of the urinary proteome of patients with sAH, AC, HD, and HC

To better understand the potential role of complement in the pathophysiology of ALD, we conducted an unbiased proteomic analysis of urine from HC and HD and patients with sAH or AC. The LC-MS data was searched against the human SwissProtKB database ^37^; 17,141 peptides were identified and mapped to 3211 proteins.

A list of 80 proteins comprising the complement system were identified based on available information ^9^; of these 80 proteins, only 51 proteins were identified in the urine proteome. Those that were significantly different between groups are highlighted in the volcano plots. Volcano plots depicted perturbations in multiple complement proteins; components of all three activation pathways both up- and down-regulated (Supplemental Figure 2).

#### Alternative pathway

Multiple complement proteins in the alternative pathway differed between the groups. CFB and CFD are activators in the alternative pathway; CFB was increased in AC and sAH, while CFD trended towards an increase in sAH (Figure 2A). CFP (Properdin) concentration was reduced in urine of patients with sAH compared with HC, HD, and AC. Interestingly, inhibitors of the alternative pathway showed mixed responses with CFI increased in AC and sAH and CFHR4 decreased in AC and sAH compared to HC and HD (Figure 2A). The FDR-adjusted p-values for all analyses of the urinary proteome are summarized in Supplemental Table 2.

**Figure 2.**
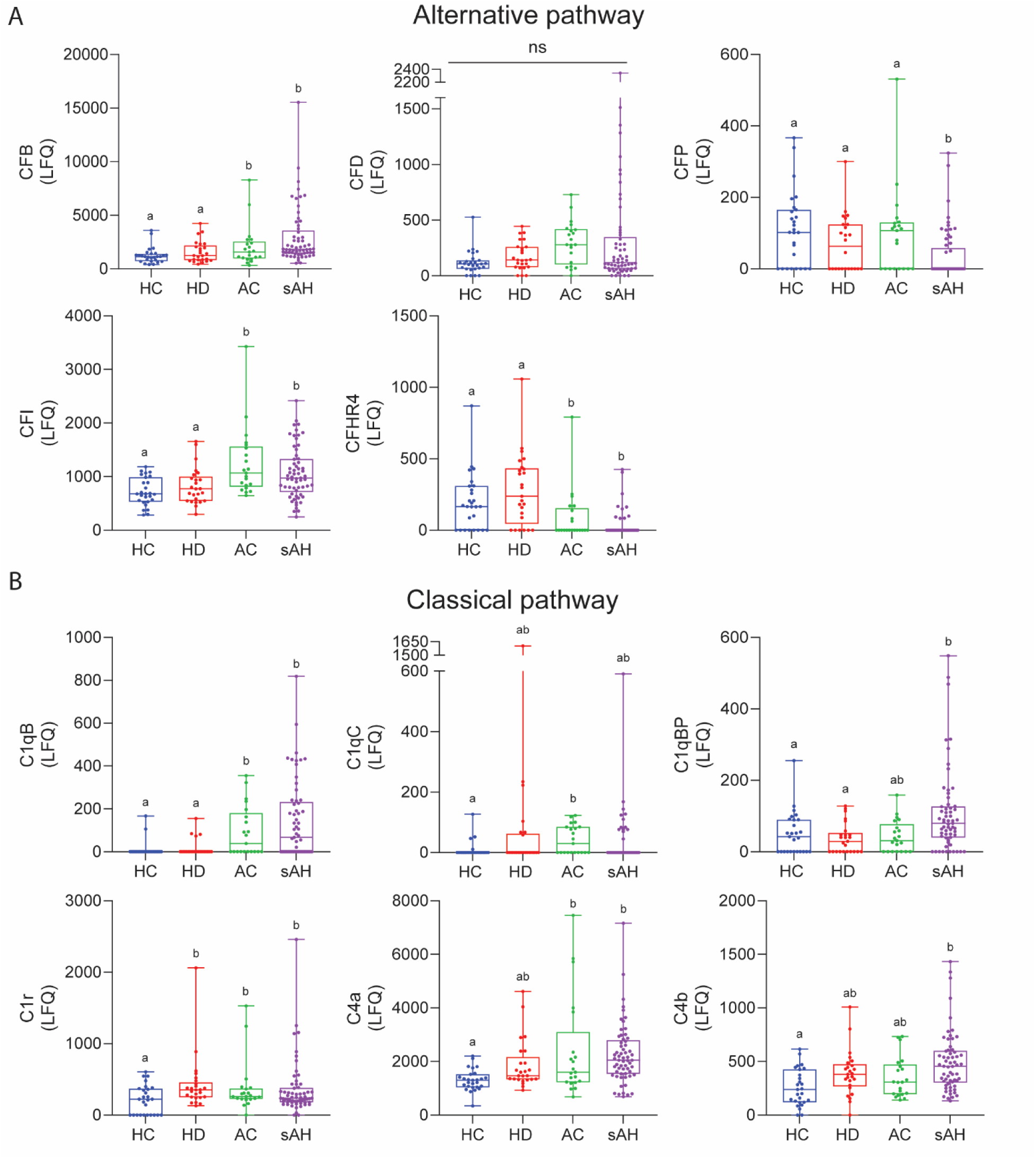
Alternative and classical pathways are perturbed in the urine of patients with sAH and AC from cohort 2. Complement proteins were detected in the urine of HC, HD, and patients with sAH and AC. A) Alternative pathway components B) Classical pathway components. Box-and-whisker plots display minimum and maximum values, lower and upper quartiles, and the median, with each individual value represented as a point superimposed on the graph. ANOVA with FDR-adjusted p-value (q<0.1), followed by Bonferroni post hoc test for individual group comparisons. Different letters mean significant differences between groups’ mean values.

#### Classical pathway

Proteins in the classical pathway showed a distinct pattern of complement perturbation between groups. Components of the C1q trimer (C1qB and C1qC), C1qBP, and C1r were generally increased in patients with ALD. Similarly, C4a and C4b, downstream effectors in the classical pathway, were also increased in patients with AC and sAH compared to HC (Figure 2B).

#### Lectin pathway

All the lectin pathway proteins identified in the urinary proteome were downregulated in patients with AC and sAH compared to HC and HD (Figure 3A). Interestingly, Umod, a protein exclusively produced in the epithelial cells of the thick ascending limb of the Loop of Henle that interacts with CL11 in the lectin pathway, was also downregulated in the urine of AC, and even further decreased in patients with sAH (Figure 3A).

**Figure 3.**
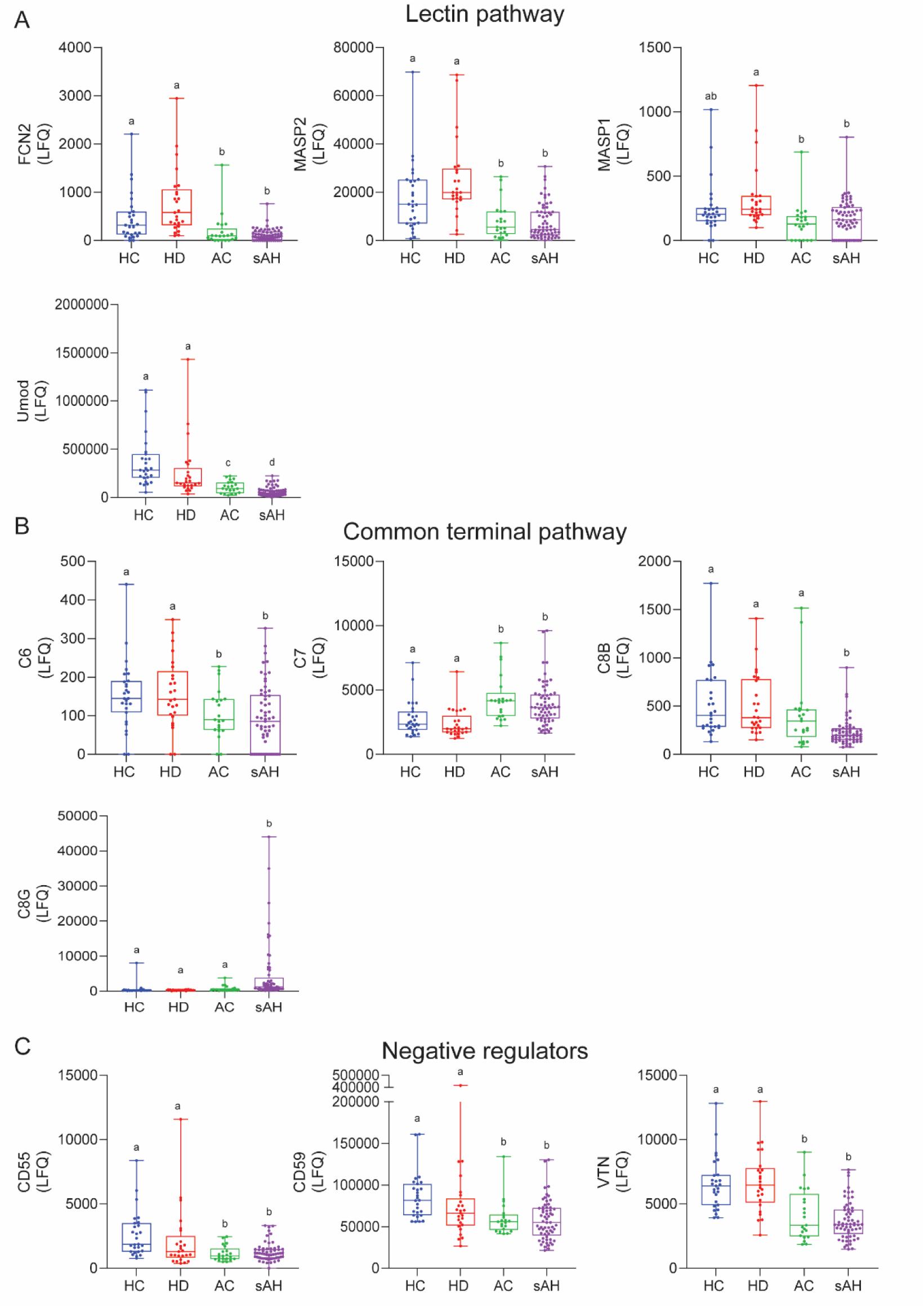
Lectin pathway, common terminal pathways and MAC inhibitors are perturbed in the urine of patients with sAH and AC from cohort 2. A) Lectin pathway activators. B) Common terminal pathway components. C) MAC formation inhibitors. Box-and-whisker plots display minimum and maximum values, lower and upper quartiles, and the median, with each individual value represented as a point superimposed on the graph. ANOVA with FDR-adjusted p-value (q<0.1), followed by Bonferroni post hoc test for individual group comparisons. Different letters mean significant differences between groups’ mean values.

#### Common terminal pathway

The common terminal pathway is responsible for assembling the membrane attack complex. There was a reduction in C6 concentration in the urine of patients with AC and sAH compared with HC and HD, whereas C7 was increased. C8 beta chain (C8B) was decreased in the urine of patients with sAH compared to HC, HD, and AC. In contrast, the C8 gamma (C8G) chain was increased (Figure 3B)

#### Negative regulators

Appropriate regulation of complement activity depends on negative regulators in targeting multiple sites within the three activation cascades to prevent overactivation. Two MAC formation inhibitors (CD55 and CD59), and VTN, were decreased in urine of patients with AC and sAH compared to HC and HD (Figure 3C). This may favor the formation of MAC and cell membrane injury, which could impact kidney health.

Taken together, these results from complement in the urine proteome demonstrate the greater impact of liver disease *per se* on the concentrations of complement in the urine of patients with AC and sAH, compared to any influence of heavy drinking.

### Complement proteins as candidate biomarkers for distinguishing sAH and AC

We next investigated if individual complement components could discriminate between patients with AC from sAH. Of all the complement proteins in the urine, Umod, C1qBP, CFP, and VSIG4, individually were the best discriminators, but only had low to moderate ability to discriminate between AC and sAH (Figure 4A). However, when the combination of all four proteins was evaluated using the leave-one-out method, the combined metric distinguished AC from sAH with an AUC of 0.78. This value was not different from the ability of MELD score, the standard tool for diagnosing sAH, to distinguish sAH from AC in this cohort, with an AUC of 0.65 (p=0.28 for t-test comparison) (Figure 4B and Table 3).

**Figure 4.**
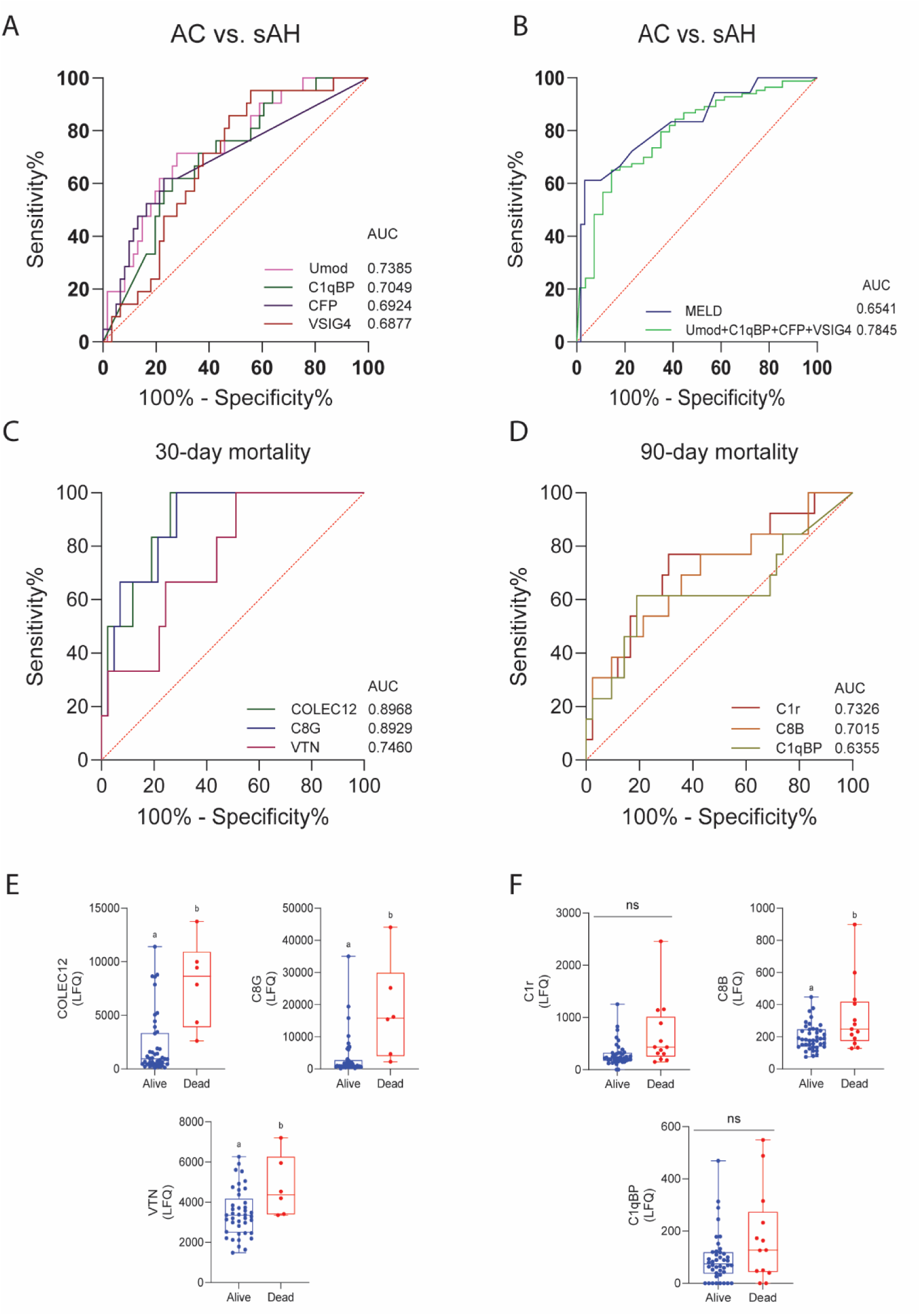
Complement proteins can distinguish sAH from AC and are associated with 30- and 90-day mortality in patients with sAH. A) ROC curve shows AUC of the top 4 proteins able to distinguish between patients with sAH and AC. B) ROC curve shows AUC of MELD and the combination of Umod, C1qBP, CFP and VSIG4 between patients with sAH and AC. 30- (C) and 90-day mortality (D) ROC curves. AUC values are in the graphs. E) Complement proteins associated with 30-day mortality comparison between patients with sAH who died and those who did not. F) Complement proteins associated with 90-day mortality comparison between patients with sAH who died and those who did not. Values were log-transformed and the comparisons were performed using student’s t-test (p<0.05).

**Table 3.** Area under the curve (AUC) values quantify the discrimination between sAH and AC and the association of complement components with 30- and 90-day mortality in patients with sAH.

|  | Protein/Predictor | AUC | Specificity (%) | Sensitivity (%) |
| --- | --- | --- | --- | --- |
| **sAH vs. AC** | Umod | 0.74 | 72 | 71 |
|  | C1qBP | 0.70 | 74 | 62 |
|  | CFP | 0.69 | 72 | 62 |
|  | VSIG4 | 0.69 | 63 | 71 |
|  | MELD<sup>†</sup> | 0.65 | 97 | 52 |
|  | Umod+C1qBP+CFP+VSIG<sup>†</sup> | 0.78 | 72 | 81 |
| **30-day mortality** | Collec12 | 0.90 | 100 | 74 |
|  | C8G | 0.89 | 100 | 71 |
|  | VTN | 0.75 | 67 | 74 |
| **90-day mortality** | C1r | 0.73 | 77 | 69 |
|  | C8B | 0.70 | 62 | 69 |
|  | C1qBP | 0.64 | 62 | 81 |

<sup>†</sup>AUC values were validated using the LOO method.

### Complement components predicted 30- and 90-day mortality in patients with sAH

The 30- and 90-day mortality rates were 9.8% and 21.3%, respectively, in patients with sAH. These mortality rates are similar to those recently reported by Dasarathy et al. ^28^, who found 13.6% and 21.5% of 30- and 90-day mortality in the complete set of patients enrolled in the AlcHepNet Observational study ^28^. Of all the complement proteins evaluated, only those that were upregulated, and not downregulated, were associated with mortality. CL12, C8G, and VTN were increased in the urine of patients and were associated with 30-day mortality in patients with sAH (Figure 4C, Table 3), while C1r, C8B, and C1qBP were associated with 90-day mortality (Figure 4D, Table 3). The pairwise analysis of the proteins with discriminatory power showed that CL12, VTN, and C8G were upregulated in the urine of patients with sAH who died within 30 days (Figure 4E). At 90 days, only C8B was upregulated in the urine of patients with sAH who died compared to those who survived (Figure 4F).

### Urinary and plasma complement concentrations do not correlate

Differences in the accumulation of complement proteins in the urine in patients with ALD could result from 1) changes in complement production in liver and other extra-renal tissues, 2) local complement synthesis in the kidney and/or 3) loss of glomerular size selectivity or impaired re-absorption of proteins in the tubules. In order to understand if perturbations in urinary complement in sAH were associated with perturbations in renal and/or extrarenal regulation of complement, we evaluated the correlations between plasma and urinary complement proteins. While contrasting individual complement proteins between plasma and urine revealed low-to-moderate Spearman correlations (Figure 5A), there were no significant correlations after FDR-adjustment (Figure 5B-I). These results suggest that it is not only the concentration of complement proteins in the plasma that is driving differences in the urinary complement proteome and that liver disease influences local renal complement production and/or handling.

**Figure 5.**
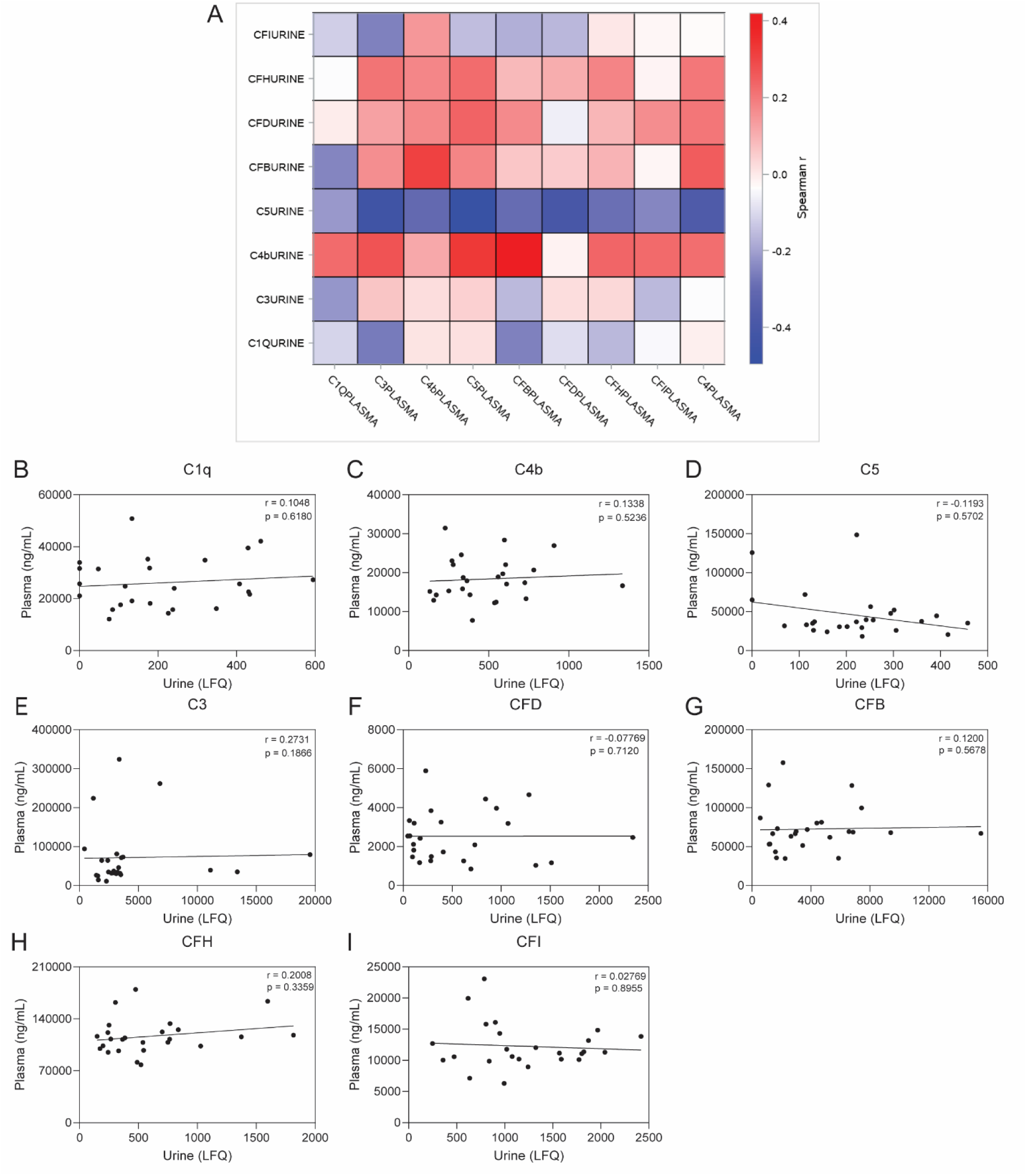
Complement correlations between urine and plasma. A) Heatmap depicting correlations between the same complement protein detected in urine and plasma from patients with sAH. Heatmap colored by Pearson’s correlation (r). Individual Spearman correlations of B) C1q; C) C4b; D) C5; E) C3; F) CFD; G) CFB; H) CFI; I) CFH.

## Discussion

Using unbiased proteomics, we characterized the urinary proteome in patients with ALD, heavy drinkers, and healthy controls. Recent publications ^9,26^ provide evidence that complement is perturbed and activated in the circulation and liver of patients with sAH. Here, we found that complement is also perturbed and activated in the urine of patients with sAH and AC. These two stages of ALD had a greater impact on complement in both urine and plasma compared to heavy drinking in the absence of liver disease.

An important finding in our study was that all three complement activation pathways were perturbed in both the plasma and urine of patients with sAH or AC. Accumulation of multiple components of both the classical and alternative pathways were increased in patients with ALD, while components of the lectin pathway were downregulated. Complement activation fragments, iC3b and C4b, were increased in the plasma and/or urine of patients with sAH, suggesting that the integrated impact of the multiple and complex perturbations in complement resulted in activation of complement in patients with ALD.

Dysregulation of the alternative pathway was characterized by upregulation of activators and downregulation of negative regulators. The alternative pathway is associated with clearance of dead cells in multiple systems, including in the livers of mice exposed to ethanol ^16^. Therefore, an overall upregulation of the alternative pathway is consistent with an increased demand for clearance of injured or dead cells that may accumulate in the liver and other organs in patients with ALD. In contrast, activators of the lectin pathway are pattern recognition receptors (PRRs) that bind to pathogens, leading to their destruction ^13,38,39^. Decreased activity of the lectin pathway could predispose patients with sAH to infections and sepsis, which are commonly associated with the disease ^40^.

Umod is a glycoprotein produced exclusively by the thick ascending limb of the Loop of Henle. Umod plays roles in kidney homeostasis and in systemic processes, including immune and inflammatory modulation, inhibition of oxidative stress, and inhibition of vascular calcification^41^. There is mounting evidence that regulation of expression of Umod impacts the outcomes of multiple diseases ^24,41–47^. The concentration of Umod was reduced in the urine of patients with sAH and AC. Additionally, the concentration of Umod in the urine of patients with sAH was significantly lower than that in patients with AC. Lower urinary Umod levels are associated with poorer prognosis for patients with AC at hospital admission ^48^. TNF-α suppresses Umod expression in epithelial cells of the thick ascending limb of the loop of Henle ^41^. It is possible that elevated TNF-α in liver disease ^49,50^ might negatively affect Umod expression and secretion into the urine, although the precise mechanism remains unclear and warrants further research. One known role of Umod is its interaction with CL11 to regulate the lectin pathway. Umod binds to CL11, an activator of the lectin pathway; this interaction mechanically inhibits the lectin pathway activation^24^. CL11 is increased in the circulation of patients with sAH (Figure 1)^9^ and may interact with Umod.

Previous work has demonstrated that expression of multiple complement proteins is perturbed in the liver of patients with sAH ^9^. While many complement proteins are produced in the liver by hepatocytes and secreted into the circulation, multiple other cells and tissues, including the kidney, also synthesize complement proteins. Local/extra-hepatic complement production can have a profound impact on local complement activation during disease ^51–56^. Indeed, kidney injury, of multiple etiologies, can lead to an increase of complement in urine, as well as complement activation in the apical membrane of the tubular epithelial cells^19^.

Interestingly, in our analysis, complement proteins were not significantly correlated between plasma and urine. These results suggest that it is not only the concentration of complement proteins in the plasma that is driving differences in the urinary complement proteome. Therefore, it is likely that liver disease influences local renal complement production, loss of glomerular size selectivity and/or the ability of tubular epithelial cells to reabsorb specific complement proteins.

The current study has several strengths and limitations. Strengths include the use of well-characterized patient cohorts enrolled in multicenter studies. The inclusion of heavy drinkers without any liver disease as a comparison with patients with ALD was essential to understand the impact of liver disease vs. alcohol consumption on complement. The use of unbiased, non-targeted proteomics is also a strength, enabling the assessment of all complement proteins in urine. A critical limitation of the study is that patients in the AlcHepNet Observational study received standard of care for their liver disease, which may vary by center and between patients, potentially influencing the observed outcomes. Additionally, liver biopsies, the gold standard for distinguishing sAH from AC, were unavailable for this study. The enrollment of mostly Caucasian/White individuals must be considered in this study, as it may not accurately reflect potential racial and ethnic differences. The samples used in this study were collected at study enrollment and may therefore not accurately reflect changes during treatment. Finally, one important limitation is that, while we did make use of leave-one-out validation strategies, an external cohort was not available for validation.

In summary, complement is perturbed in the urine of patients with ALD, including sAH and AC, and this disturbance is likely due to complex cross-talk between the liver and kidney in patients with ALD. Importantly, when multiple complement proteins are combined, they are as powerful as MELD score to distinguish patients with AC from sAH. The findings of this work provide important insights into the mechanisms underlying liver-kidney crosstalk and how liver disease can affect kidney health in patients with ALD.

## Data availability

Data is available in the Supporting Data Values Excel file.

## Author contributions

LP and LEN conceptualized the work. AlcHepNet consortium and Northern Ohio Alcohol Center (SD, JD, NW, DS) collected samples. LP and MT conducted assays. RM performed all the bioinformatics analysis. LP, LEN, VP, DMR, and RM analyzed the data. LP wrote the first draft of the manuscript. LP and LEN wrote and edited the manuscript. All authors reviewed and approved the manuscript.

