## Supplemental figures for "Urine proteomic profiling at admission reveals complement biomarkers linked to alcohol-associated liver disease"

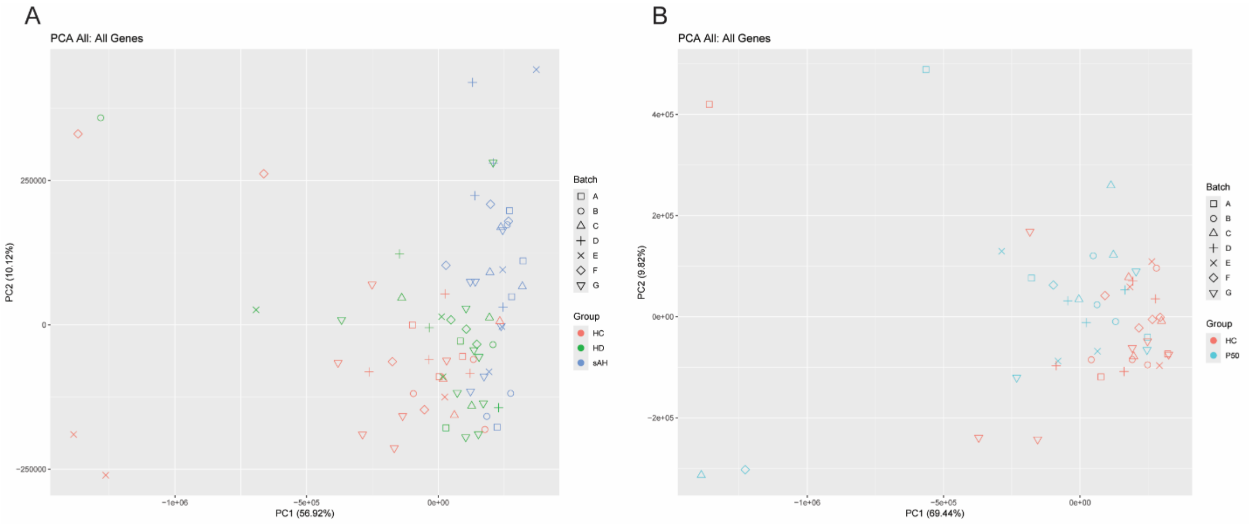


**Supplemental figure 1. PCA plot for all the proteins found in the urinary proteome.** Samples were analyzed by liver disease and batch to exclude batch effects for A) HC, HD and sAH; and B) HC and AC.

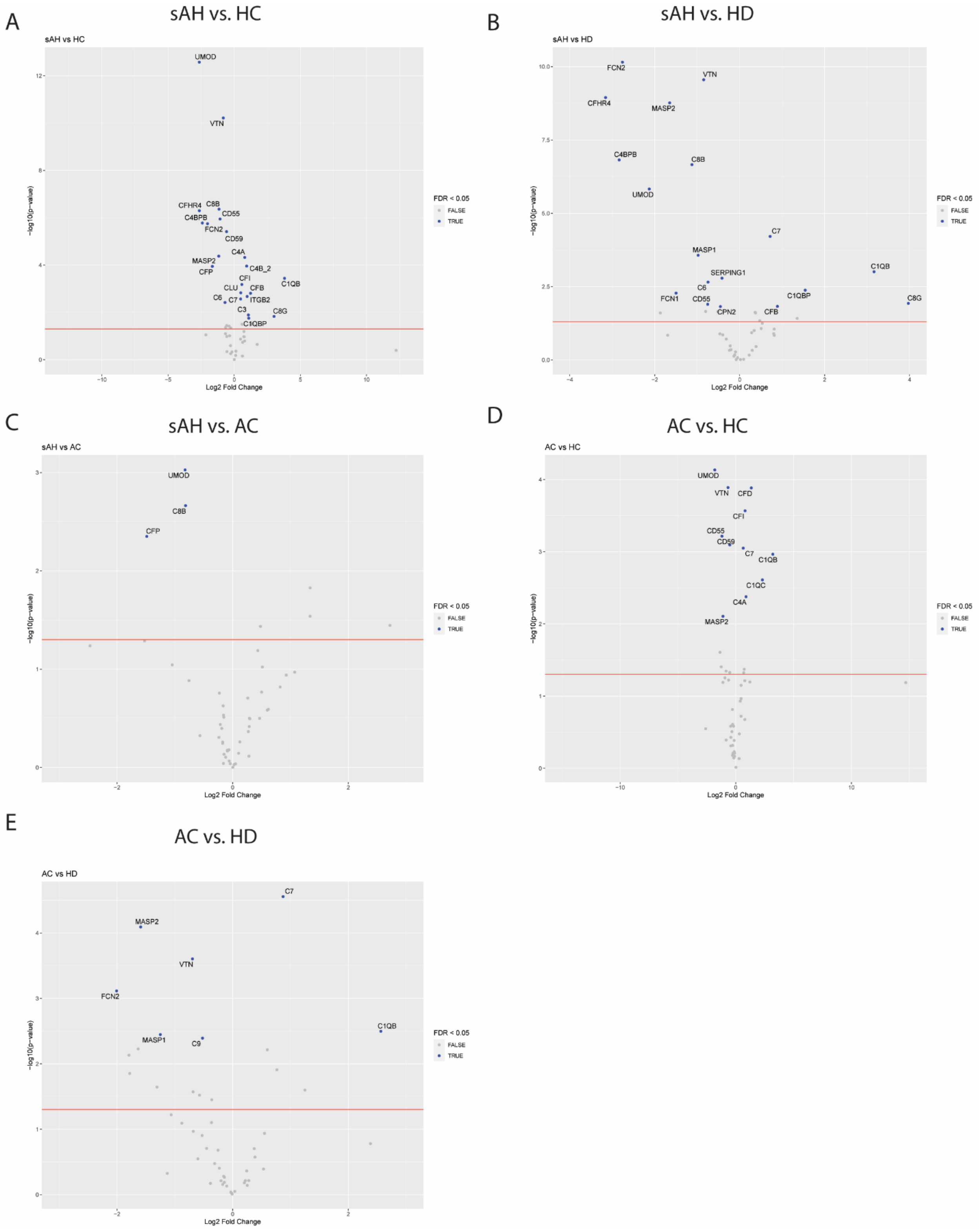


**Supplemental Figure 2. Complement protein expression in the urine of patients with sAH and AC, HC and HD individuals.** The volcano plots depict pairwise comparisons between A) sAH vs. HC; B) sAH vs. HD; C) AC vs. sAH; D) AC vs. HC; E) AC vs. HD.
