## Supplemental tables for "Urine proteomic profiling at admission reveals complement biomarkers linked to alcohol-associated liver disease"

Table 1. FDR-adjusted p-values for the comparisons between HC, HD and sAH in cohort 1.

| Variables | p-value | FDR adjusted p-value |
|---|---|---|
| C4b | <0.0001 | 0.00019 |
| CFB | <0.0001 | 0.00019 |
| CFD | <0.0001 | 0.00019 |
| FH | <0.0001 | 0.00019 |
| FI | <0.0001 | 0.00019 |
| iC3B | <0.0001 | 0.00019 |
| CL11 | <0.0001 | 0.00019 |
| C5E | 0.0293 | 0.04761 |
| C1q | 0.0521 | 0.06773 |
| MBL | 0.0482 | 0.06773 |
| sC5B9 | 0.1454 | 0.16759 |
| C3 | 0.1547 | 0.16759 |
| C5 | 0.1755 | 0.17550 |

Table 2. FDR-adjusted values for the comparisons between HC, HD, AC, and sAH in cohort 2.

| Variables | p-value | FDR adjusted p-value |
|---|---|---|
| VTN | <0.0001 | 0.00013 |
| CD59 | <0.0001 | 0.00013 |
| CLU | <0.0001 | 0.00013 |
| C7 | <0.0001 | 0.00020 |
| C8B | <0.0001 | 0.00020 |
| C8G | <0.0001 | 0.00020 |
| CFB | <0.0001 | 0.00030 |
| CFHR4 | <0.0001 | 0.00030 |
| CFI | <0.0001 | 0.00030 |
| C1QB | <0.0001 | 0.00037 |
| C1R | <0.0001 | 0.00037 |
| C4BPB | <0.0001 | 0.00037 |
| FCN2 | <0.0001 | 0.00045 |
| UMOD | <0.0001 | 0.00045 |
| C4A | 0.0002 | 0.00055 |
| MASP2 | 0.0002 | 0.00060 |
| CFP | 0.0003 | 0.00068 |
| MASP1 | 0.0025 | 0.00563 |
| C4B | 0.0081 | 0.01782 |
| C6 | 0.0147 | 0.02205 |
| C1QBP | 0.0153 | 0.02805 |
| C1QC | 0.0190 | 0.02986 |
| C4BPA | 0.0282 | 0.03878 |
| FCN1 | 0.0233 | 0.04194 |
| CD55 | 0.0348 | 0.06264 |
| C1S | 0.0605 | 0.07394 |
| CL10 | 0.0577 | 0.08655 |
| CFH | 0.0581 | 0.08715 |
| CD93 | 0.0869 | 0.09559 |
| CL12 | 0.0834 | 0.10723 |
| FCN3 | 0.1110 | 0.12488 |
| C2 | 0.2203 | 0.22030 |
| SERPING1 | 0.2542 | 0.25420 |
| CFD | 0.3257 | 0.41876 |
| CD209 | 0.6383 | 0.63830 |
| C8A | 0.6414 | 0.66060 |
| C5 | 0.6606 | 0.66060 |
| VSIG4 | 0.6714 | 0.75533 |
| CFHR2 | 0.8507 | 0.85070 |

Supplemental Table 3: "AlcHepNet Clinical Center Investigators" Roster for Authorship

| University/Center | Authors |
|---|---|
| **Indiana University School of Medicine, Indianapolis, IN** | Naga Chalasani, MD; Ojo Olawale, MD; Lauren Nephew, MD; Raj Vuppalanchi, MD; Niha Samala, MD; Lindsey Yoder, PA; Suthat Liangpunsakul, MD, MPH, Caitlin Snoeberger, RN |
| **Mayo Clinic, Rochester, Minnesota** | Vijay H. Shah, MD; Douglas A. Simonetto, MD; Patrick Kamath, MD; Victor Karpyak, MD; Amy Olofson, RN |
| **Cleveland Clinic, Cleveland, Ohio** | Srinivasan Dasarathy, MD; Nicole Welch, MD; Annette Bellar; Amy Attaway, MD; Jaividhya Dasarathy, MD (MetroHealth, Cleveland, Ohio); Ashley Growley; David Streem, MD; Laura E. Nagy, PhD |
| **University of Texas Southwestern, Dallas, Texas** | Mack C. Mitchell, MD; Thomas Kerr, MD; Thomas Cotter, MD; Jacquelyn O'Leary, MD; Sherwood Brown, MD |
| **Virginia Commonwealth University, Richmond, Virginia** | Arun J. Sanyal, MD, MBBS; Sara O'Connor, RN; Velimir Luketic, MD; Amon Asgharpour, MD; Juan Pablo Arab MD; Richard K Sterling MD MSc; F Gerard Moeller, MD |
| **University of Louisville, Louisville, Kentucky** | Craig J. McClain, MD; Ashwani K. Singal, MD; Christopher Stewart, MD; Vatsalya Vatsalya, MD, PhD; (Matt Cave, MD; Luis Marsano, MD; Ashutosh Barve, MD, PhD; and Jane Frimodig, PhD- Collaborators) |
| **Data Coordinating Center at Indiana University** | Wanzhu Tu, PhD; Samer Gawrieh, MD; Qing Tang, MS; Carla Kettler, MS; Jing Su, PhD; Yang Li, PhD; Savannah Yarnelle; Yunpeng Yu, MS; Tae-Hwi L. Schwantes-An, PhD |
